# Gene knock out of honey bee trypanosomatid parasite, *Lotmaria passim*, by CRISPR/Cas9 system

**DOI:** 10.1101/478198

**Authors:** Qiushi Liu, Tatsuhiko Kadowaki

## Abstract

Two trypanosomatid species, *Lotmaria passim* and *Crithidia mellificae,* have been shown to parasitize honey bees to date. *L. passim* appears to be more prevalent than *C. mellificae* and specifically infects the honey bee hindgut. Although the genomic DNA has been sequenced, the effects of infection on honey bee health and colony are poorly understood. To identify the genes that are important for infecting honey bees and to understand their functions, we applied the CRISPR/Cas9 system to establish a method to manipulate *L. passim* genes. By electroporation of plasmid DNA and subsequent selection by antibiotics, we first established an *L. passim* clone expressing tdTomato, GFP, or Cas9. We also successfully knocked out the endogenous miltefosine transporter and tyrosine amino transferase genes by replacement with antibiotics (hygromycin) resistant gene using the CRISPR/Cas9-induced homology-directed repair pathway. The *L. passim* clones expressing fluorescent markers, as well as the simple method for knocking out specific genes, could become useful approaches to understand the underlying mechanisms of honey bee-trypanosomatid parasite interactions.

## Introduction

The honey bee (*Apis mellifera*) plays important roles in agricultural crop production and ecosystem conservation across the globe (Klein et al., 2007;Aizen et al., 2008;Potts et al., 2010). However, a decline in managed honey bee colonies has been observed in North America, Europe, and a part of Asia since 2006. The underlying reasons for the large-scale colony losses are complex, but can be divided into several different categories: inadequate food supplies, anthropogenic chemicals, and exposure to various pathogens/parasites (Goulson et al., 2015). There are diverse honey bee pathogens/parasites, such as viruses, bacteria, fungi, protozoans, and mites (Evans and Schwarz, 2011). Among the protozoans, two *Trypanosomatidae, Lotmaria passim* and *Crithidia mellificae*, were shown to infect honey bees. *C. mellificae* was first identified in Australia in 1967 (Langridge and McGhee, 1967). Later, in 2015, another novel trypanosomatid parasite infecting honey bees was discovered and named *L. passim* (Schwarz et al., 2015). *L. passim* was found to be more prevalent than *C. mellificae* (Schmid-Hempel and Tognazzo, 2010; Morimoto et al., 2013; Cepero et al., 2014; Ravoet et al., 2014; Cersini et al., 2015; Ravoet et al., 2015; Schwarz et al., 2015; Arismendi et al., 2016; Cavigli et al., 2016; Stevanovic et al., 2016; Vavilova et al., 2017; Regan et al., 2018) and fewer honey bee colonies were reported to be infected by *C. mellificae* (Ravoet et al., 2015). Thus, *L. passim* rather than *C. mellificae* is likely to be associated with the previously reported winter mortality of honey bee colonies (Ravoet et al., 2013). However, the effects of *L. passim* infection on honey bee health and colonies is poorly understood. *L. passim* specifically infects the honey bee hindgut and triggers the expression of antimicrobial peptide (AMP) genes, such as *Defensin 1* and *Abaecin*. In addition, *L. passim* stimulates the increased expression of genes encoding several downstream components of immune pathways (honey bee orthologs of Imd and Dscam) (Schwarz and Evans, 2013). Although the genome of *L. passim* has been sequenced (Runckel et al., 2014), both *C. mellificae* and *L. passim* have not been fully investigated to date.

*Crithidia bombi*, a trypanosomatid parasite of the bumble bee, has been well characterized. *C. bombi* infection dramatically reduces colony-founding success, male production, and colony size (Brown et al., 2003). Furthermore, *C. bombi* infection was also reported to impair the ability of bumble bees to utilize floral information (Gegear et al., 2006). *C. bombi* infection induces expression of several immune-related genes: *MyD88*, *Relish*, *Thioester-containing protein 7* (Schlüns et al., 2010), as well as AMP such as *Abaecin*, *Defensin* and *Hymenoptaecin* in bumble bees (Riddell et al., 2011; Riddell et al., 2014). The genomes of *C. bombi* and *Crithidia expoeki* were recently sequenced and the sequences revealed signs of concerted evolution of genes potentially important for interaction with the host (Schmid-Hempel et al., 2018).

Recently, a new method based on the CRISPR/Cas9 system has become widely used for genome editing. It has also been applied to edit the genomes of various trypanosomatid parasites: *Trypanosoma cruzi* (Lander et al., 2015a; Peng et al., 2015; Lander et al., 2016; Lander et al., 2017), *Trypanosoma brucei* (Beneke et al., 2017; Rico et al., 2018), *Leishmania major* (Sollelis et al., 2015; Beneke et al., 2017), *Leishmania donovani* (Zhang and Matlashewski, 2015; Martel et al., 2017; Zhang et al., 2017), and *Leishmania mexicana* (Beneke et al., 2017). Although the nonhomologous end-joining (NHEJ) pathway appears to be absent in trypanosomatid parasites (Passos-Silva et al., 2010), the endogenous genes were successfully knocked out both by the microhomology-mediated end joining (MMEJ) and homology-directed repair (HDR) pathways, in order to repair Cas9-induced double-strand DNA breaks (DSBs). In this study, we first generated *L. passim* clones expressing fluorescent markers and then attempted to use CRISPR/Cas9 for genome editing. We will discuss how these approaches can be used to better understand honey bee-trypanosomatid parasite interactions.

## Materials and Methods

### Culture of *L. passim*

*L. passim* strain SF (PRA-403) was obtained from the American Type Culture Collection (ATCC) and cultured in the modified FPFB medium (Salathe et al., 2012). The parasite lines with fluorescent markers were cultured in the modified FPFB medium containing 5 μg/mL blasticidin (InvivoGen). The gene knocked-out parasite lines were maintained in the presence of 5 μg/mL blasticidin, 10 μg/mL hygromycin (SIGMA), 50 μg/mL G418 (SIGMA). To monitor the growth rate, *L. passim* was first inoculated at 5 x 10^5^/mL, and then the number of parasites during the culture was measured by CASY^®^ Cell Counter together with Analyzer System Model TT (OMNI Life Science).

### Electroporation of *L. passim* followed by the single clone selection

Actively growing *L. passim* was collected, washed twice, and resuspended in 0.4 mL of Cytomix buffer (20 mM KCl, 0.15 mM CaCl_2_, 10 mM K_2_HPO_4_, 25 mM HEPES and 5 mM MgCl_2_, pH 7.6). 2 x 10^7^ of *L. passim* were electroporated with 10 μg of plasmid DNA: pTrex-Neo-tdTomato (Canavaci et al., 2010), pTrex-n-eGFP (Peng et al., 2015), or pTrex-b-NLS-hSpCas9 (Peng et al., 2015) using a Gene Pulser X cell electroporator (Bio-Rad). For co-transfection of sgRNA expression vector and donor DNA, 10 μg of the plasmid DNA and 25 μg of linearized donor DNA were electroporated. The 2 mm gap cuvettes were chilled on ice for 15 min before electroporation. The voltage, capacitance, and resistance were set at 1500 V, 25 μF, and Infinity, respectively for all experiments. The electroporation was repeated twice and there was 10 sec interval between each pulse. For single clone selection, the electroporated parasites growing in the medium with appropriate antibiotics were diluted and spread on 2.5 % agarose containing the modified FPFB medium and the antibiotics at 25 °C. The individual parasite colonies were picked and expanded in 12-well plate. We used serial dilution for the single clone selection to compare gene knock-out by CRISPR/Cas9-induced HDR and homologous recombination.

### Western blot analysis

The parasites expressing Cas9 were directly suspended with the sample buffer for SDS-PAGE and heated for 3.5 min. The cell lysates were separated by two 8 % SDS-PAGE gels and then the proteins in one gel were transferred to a PVDF membrane. Another gel was stained by coomassie brilliant blue as the loading control. The membrane was blocked with 5 % BSA/TBST and then incubated with rabbit anti-FLAG antibody (Sigma-Aldrich, 100-fold dilution) at 4 °C overnight. After washing the membrane three times with TBST, it was incubated with IRDye^®^ 680RD anti-rabbit secondary antibody (10, 000-fold dilution) in 5 % skim milk/TBST at room temperature for 1 h. The membrane was washed as above and scanned/analyzed by Odyssey^®^ scanner (LI-COR Biosciences).

### Gene knock-out by CRISPR/Cas9-induced HDR

*L. passim* miltefosine transporter (LpMT) and tyrosine amino transferase (LpTAT) sgRNA sequences were designed using a custom sgRNA design tool (http://grna.ctegd.uga.edu) (Peng et al., 2015). These two sgRNA sequences were cloned into pSPneogRNAH vector (Zhang and Matlashewski, 2015). The donor DNA for *LpTAT* gene was constructed by fusion PCR of three DNA fragments, 5’ (438 bp) and 3’ UTRs (500 bp) of *LpTAT* and the ORF of hygromycin B phosphotransferase gene derived from pCsV1300 (Park et al., 2013). Similarly, the donor DNA for *LpMT* was prepared as above except the 5’ UTR (540 bp) and the part of ORF downstream of the sgRNA target site (500 bp) were used for the fusion PCR. The fusion PCR products were cloned into EcoRV site of pBluescript II SK(+) and the linearized plasmid DNA by HindIII was used for electroporation as mentioned above. After co-transfection, the antibiotics resistant clones were selected as above.

### Genomic PCR

Genomic DNA was extracted from the parasites using DNAiso (TAKARA) and PCR was carried out using KOD FX polymerase (TOYOBO) and the specific primers shown in the Figures and Supplementary Table 1. Some of the PCR products were gel purified and directly sequenced.

### Detection of *LpMT* and *LpTAT* mRNAs by RT-PCR

Total RNA was extracted from wild type, *LpMT* and *LpTAT* heterozygous and homozygous KO parasites using TRIzol reagent (SIGMA) and treated by 1U of RNase-free DNase (Promega) at 37 °C for 30 min. 0.2 μg of total RNA was reverse transcribed by ReverTra Ace (TOYOBO) and random primer followed by PCR with KOD FX polymerase (TOYOBO) and gene specific primers listed in Supplementary Table 1.

## Results

### Generation of *L. passim* expressing fluorescent marker or Cas9

To test if we could generate an *L. passim* clone stably expressing an exogenous protein, *L. passim* was electroporated with plasmid DNA carrying tdTomato and the neomycin resistance gene (*Neo*) driven by the *Trypanasoma cruzi* rRNA promoter (pTrex-Neo-tdTomato) (Canavaci et al., 2010), followed by G418 selection. Several G418-resistant clones were isolated from an agar plate and expanded. As shown in Fig. 1, all the parasite cells expressed tdTomato; however, most of them lost the expression after 14 weeks in culture (25 passages) without G418. These results demonstrate that the electroporated plasmid DNA existed as episomal DNA, without integrating into the parasite’s chromosomal DNA. The efficiency of transient transfection was low (up to 1.21 %), indicating that the selection of stable transfectants by antibiotics is essential. We then introduced plasmid DNA containing Cas9 and the blasticidin resistance gene (*Bsd*) driven by the *T. cruzi* rRNA promoter (pTrex-b-NLS-hSpCas9) (Peng et al., 2015) into *L. passim,* followed by blasticidin selection. Expression of the FLAG-tagged Cas9 protein was confirmed by western blotting (Fig. 2A) and the growth rate of the Cas9-expressing clone in the presence of blasticidin was comparable to that of the wild type without blasticidin (Fig. 2B).

**Figure 1.**
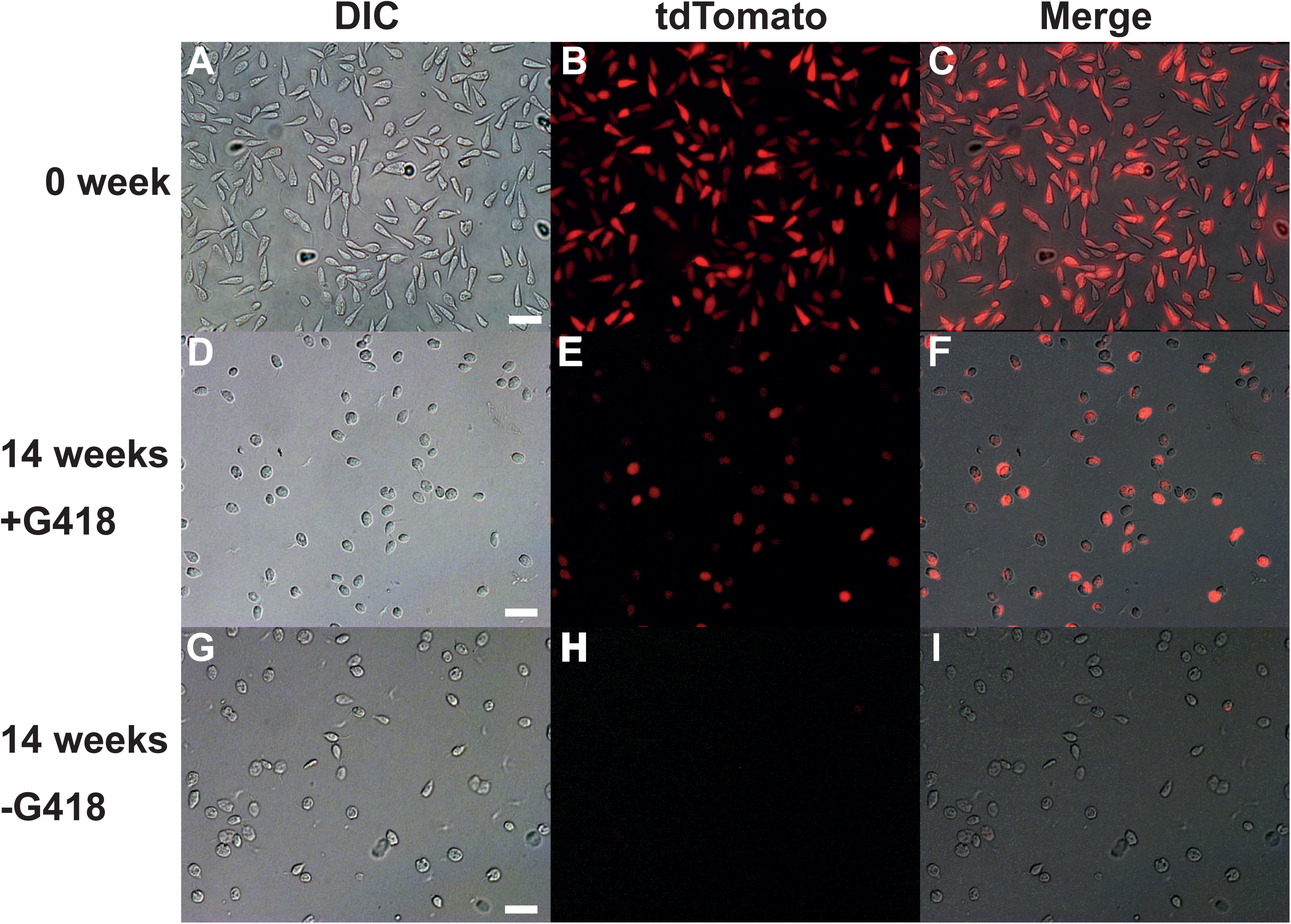
*L. passim* clone expressing tdTomato. All parasites are labelled by red fluorescence in the selected *L. passim* clone expressing tdTomato (0 week, A-C). The parasites keep expressing tdTomato when cultured for 14 weeks with G418 (14 weeks +G418, D-F) but not without G418 (14 weeks-G418, G-I). White bar = 30 μm.

**Figure 2.**
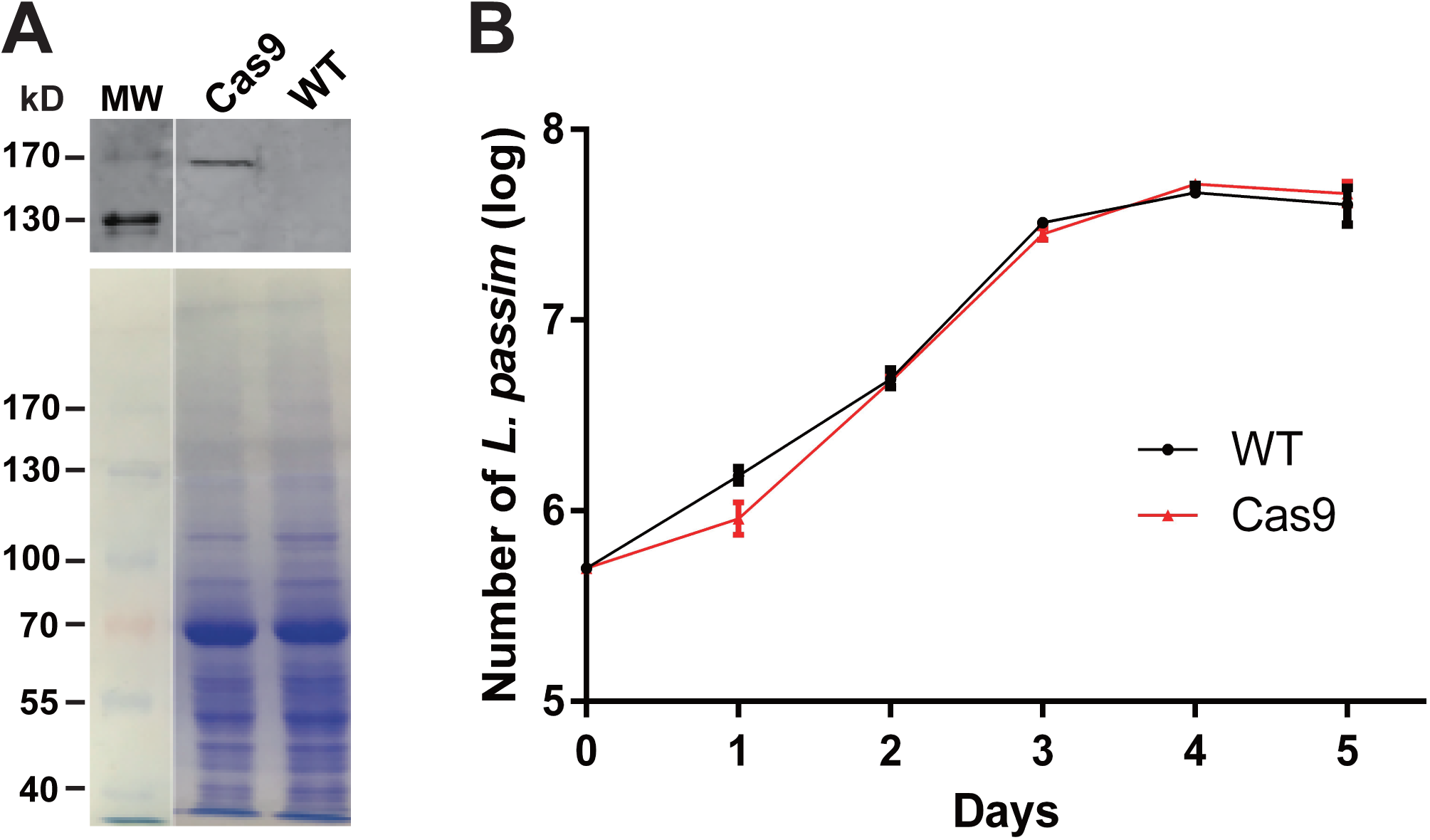
Cas9 protein expression and growth of L. passim clone expressing Cas9. **(A)** The cell lysates of wild type (WT) and FLAG-tagged Cas9 expressing (Cas9) *L. passim* were analyzed by western blot using anti-FLAG antibody (upper panel). The same SDS-PAGE gel was stained by coomassie brilliant blue as the loading control (lower panel). The size (kD) of protein molecular weight marker (MW) is at the left. **(B)** Growth of WT (black) and Cas9 (red) *L. passim* in the modified FPFB medium. The experiment was repeated three times and the error bars indicate the standard deviations.

### Knockout of miltefosine transporter and tyrosine aminotransferase genes in L. passim by Cas9-induced HDR

We first targeted the *L. passim* miltefosine transporter (*LpMT*) gene for knockout by CRISPR/Cas9-induced HDR since it is not an essential gene, and has been successfully knocked out in *Leishmania donovani* previously (Zhang and Matlashewski, 2015; Zhang et al., 2017). We transfected Cas9-expressing *L. passim* with plasmid DNA that drives the expression of *LpMT*-specific sgRNA and *Neo* under the *L. donovani* rRNA promoter (Zhang and Matlashewski, 2015), and a donor DNA. The donor DNA contained the hygromycin resistance gene (*Hyg*) flanked by 5’UTR (left arm) and a part of ORF downstream of the sgRNA targeting site of *LpMT* (right arm). After the transfection, we selected and expanded the blasticidin-, G418-, and hygromycin-resistant clones. As shown in Fig. 3A, all the antibiotic resistant clones had the *LpMT* knockout allele mediated by *Hyg* insertion through homologous recombination; however, three out of 11 clones also contained the wild type allele, suggesting that they were heterozygous. To confirm the knockout of *LpMT*, we examined the mRNA expression in wild type, heterozygous, and homozygous knockout parasites by RT-PCR. *LpMT* mRNA was absent in the homozygous knockout strain as shown in Fig. 3B. We also successfully knocked out tyrosine aminotransferase (*LpTAT*) using Cas9-induced HDR as described above (Fig. 4). *LpTAT* is one of the *L. passim* genes that becomes upregulated upon infection of the honey bee hindgut. As shown in Fig. 5, the growth rate of *LpMT* or *LpTAT* knockout parasites in culture medium was lower than that of wild type, suggesting that these two genes are not essential for the parasites’ survival, but are necessary to support the optimal growth of *L. passim*.

**Figure 3.**
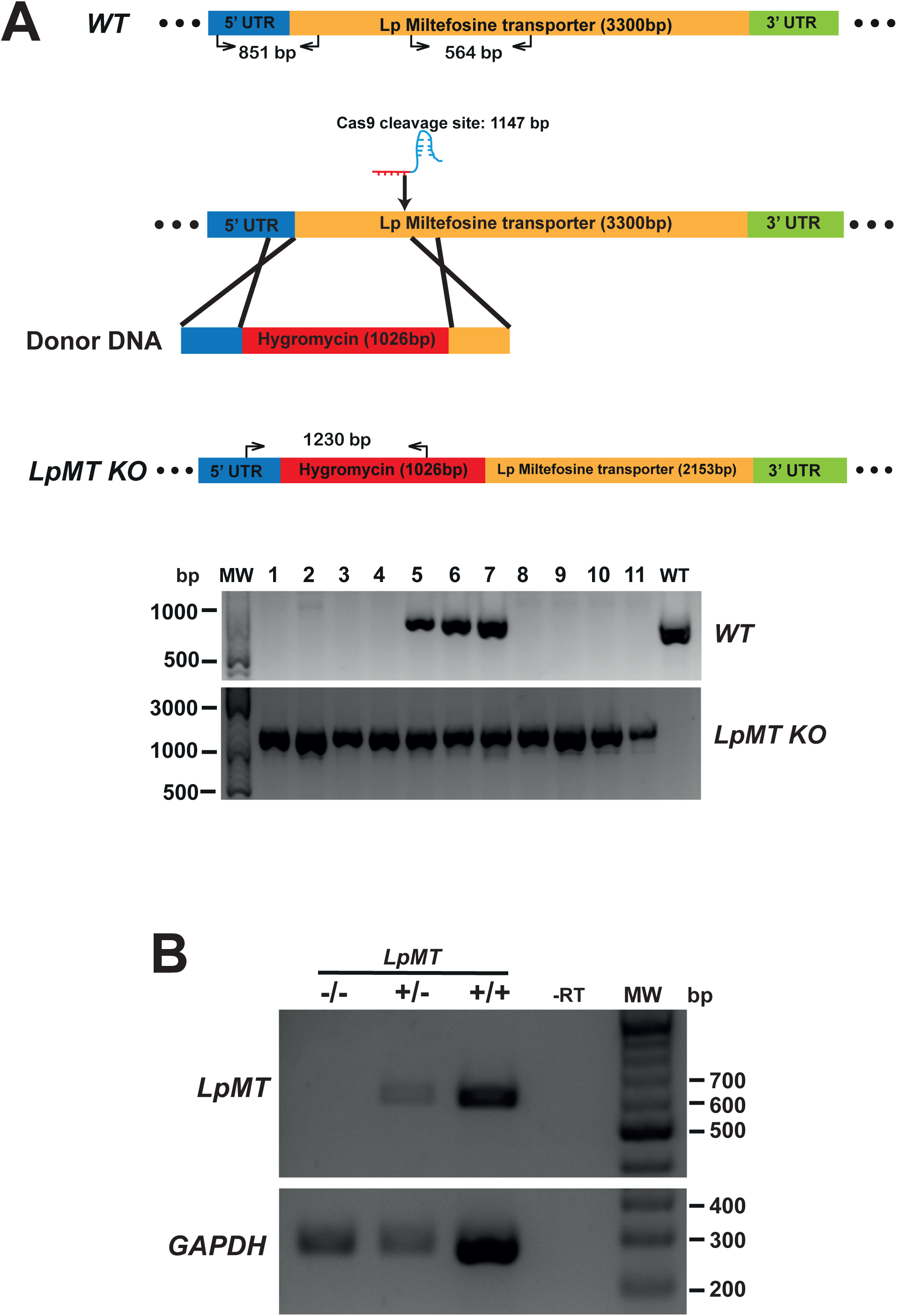
Generation of miltefosine transporter knocked out *L. passim* by CRISPR/Cas9-induced homology directed repair. **(A)** Schematic representation of the strategy used to generate *L. passim miltefosine transporter* knocked out (*LpMT KO*) parasite by CRISPR/Cas9-induced homology directed repair (HDR). The positions of primers to detect wild type (*WT*) and *KO* alleles of *LpMT* are shown with the expected sizes of PCR amplicons. 5’ and 3’ untranslated regions (UTR) as well as open reading frame (ORF) of *LpMT* are shown in blue, green, and yellow, respectively. Donor DNA contains a hygromycin resistance gene (red) flanked by the parts of 5’UTR and ORF of *LpMT*. The putative cleavage site by Cas9 is at 1,147 bp from the start codon of *LpMT*. 11 antibiotics (blasticidin, G418, and hygromycin) resistant clones (1-11) together with wild type *L. passim* were analyzed by genomic PCR to detect *WT* (851 bp amplicon) and *KO* (1230 bp amplicon) alleles of *LpMT*. The position of 500, 1000, and 3000 bp DNA molecular weight marker (MW) is shown at the left. **(B)** Detection of *LpMT* and *GAPDH* mRNAs in *LpMT* heterozygous (+/-) and homozygous (-/-) knocked-out together with wild type *L. passim* (+/+) by RT-PCR. The expected sizes of RT-PCR amplicons for *LpMT* and *GAPDH* are 564 bp and 279 bp, respectively. The negative control was run using water as the template (-RT) for RT-PCR. The position of 200-700 bp DNA molecular weight marker (MW) is shown at the right.

**Figure 4.**
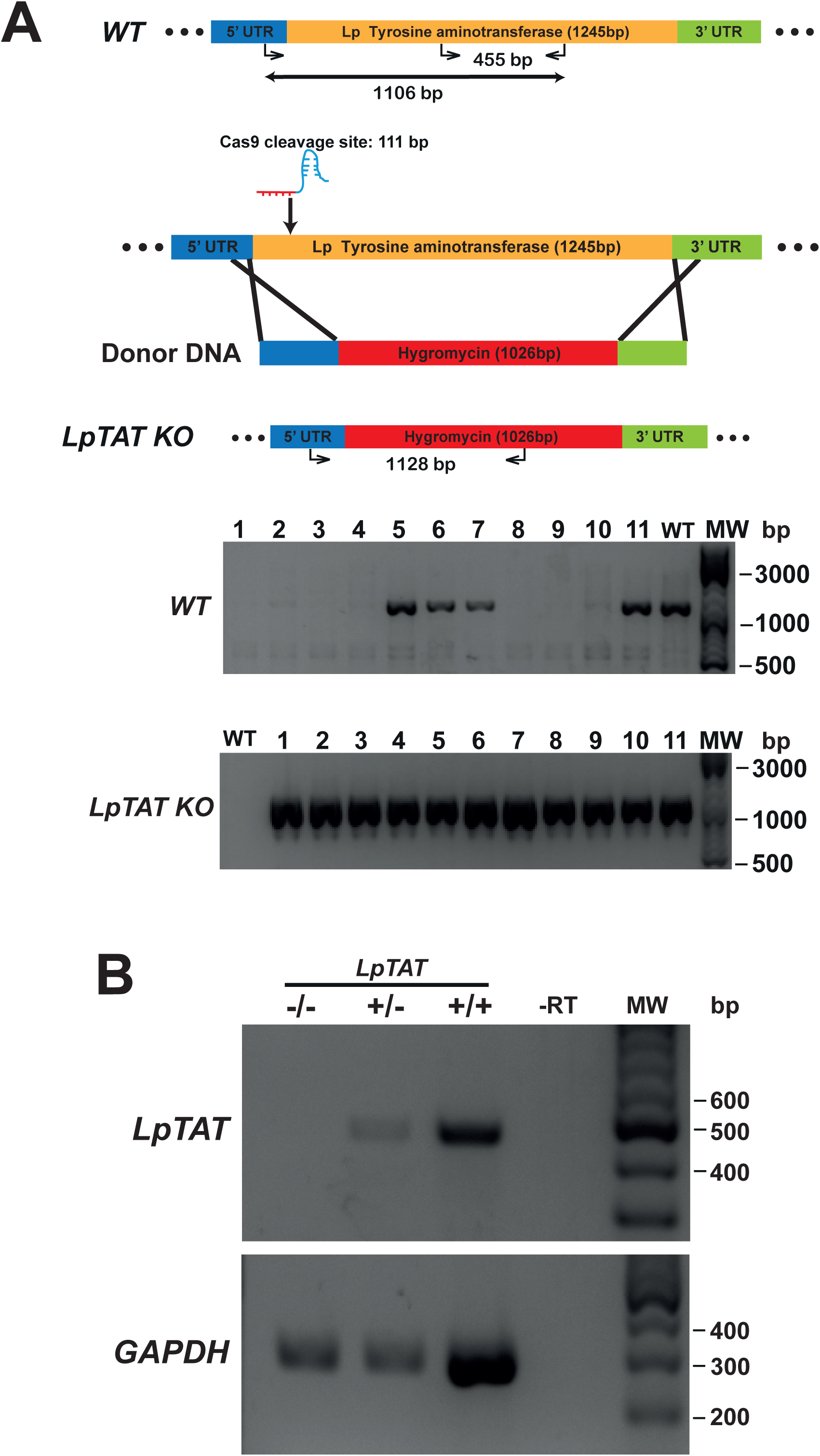
Generation of tyrosine amino transferase knocked out *L. passim* by CRISPR/Cas9-induced HDR. **(A)** Schematic representation of the strategy used to generate *L. passim tyrosine amino transferase* knocked out (*LpTAT KO*) parasite by CRISPR/Cas9-induced HDR is shown as in Fig. 3A. Donor DNA contains a hygromycin resistance gene (red) flanked by the parts of 5’ and 3’UTRs of *LpTAT*. The putative cleavage site by Cas9 is at 111 bp from the start codon of *LpTAT*. 11 antibiotics (blasticidin, G418, and hygromycin) resistant clones (1-11) together with wild type *L. passim* were analyzed by genomic PCR to detect *WT* (1106 bp amplicon) and *KO* (1128 bp amplicon) alleles of *LpTAT*. The position of 500, 1000, and 3000 bp DNA molecular weight marker (MW) is shown at the right. **(B)** Detection of *LpTAT* and *GAPDH* mRNAs in *LpTAT* heterozygous (+/-) and homozygous (-/-) knocked-out together with wild type *L. passim* (+/+) by RT-PCR. The expected sizes of RT-PCR amplicons for *LpTAT* and *GAPDH* are 455 bp and 279 bp, respectively. The negative control was run using water as the template (-RT) for RT-PCR. The position of 200-600 bp DNA molecular weight marker (MW) is shown at the right.

**Figure 5.**
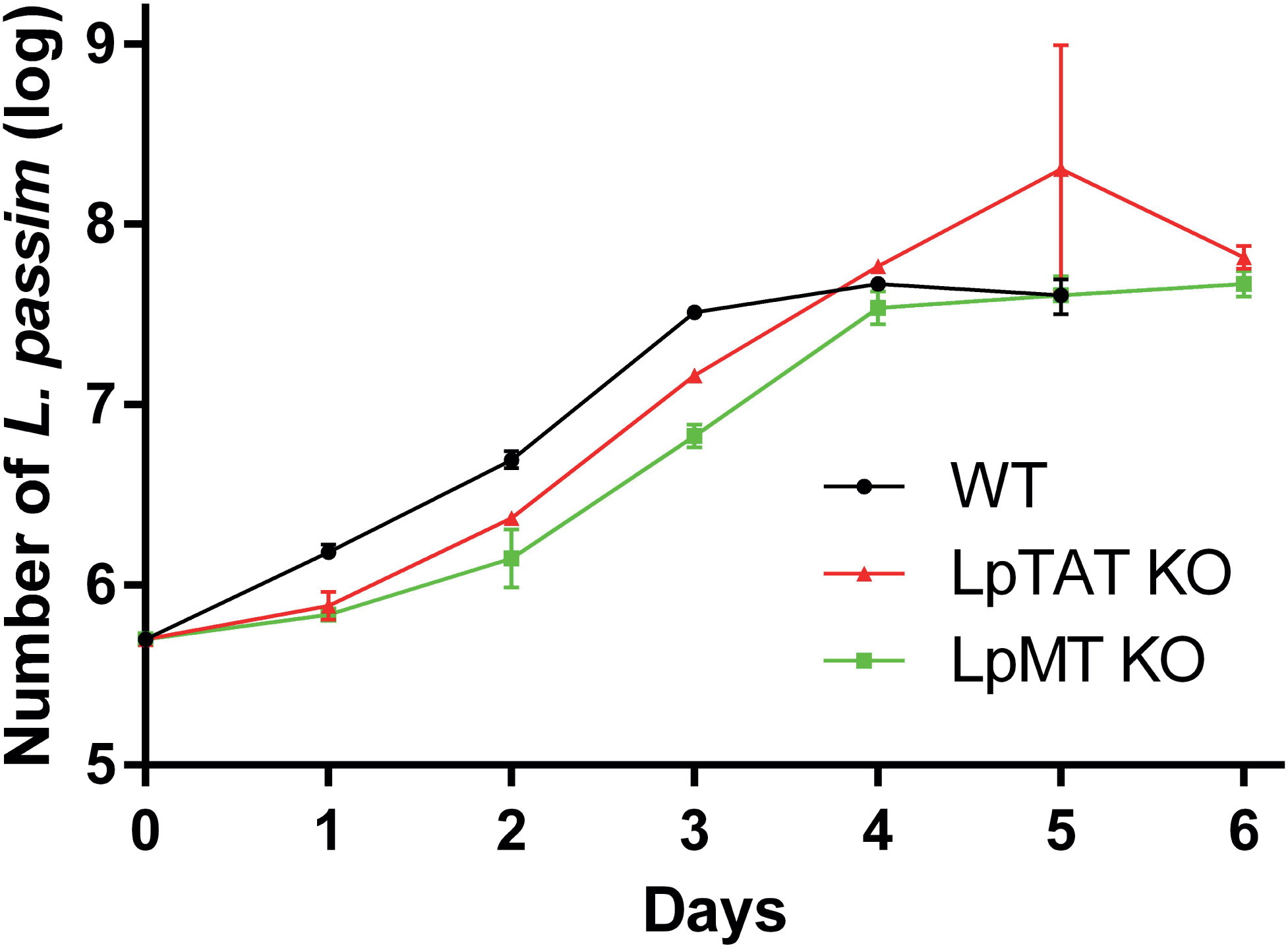
Growth of *LpTAT* and *LpMT* homozygous knocked out together with wild type *L. passim*. Growth of *LpTAT* (red) and *LpMT* (green) homozygous knocked out together with wild type (black) *L. passim* in the modified FPFB medium was measured. The experiment was repeated three times and the error bars indicate the standard deviations. Two different clones were tested for each homozygous knocked out parasite.

To confirm that replacing the endogenous gene with the donor DNA containing *Hyg* was mediated by DNA break-induced HDR rather than homologous recombination, we repeated the knocking out of *LpMT* as described above, together with transfecting wild type *L. passim* with the donor DNA only. After incubation in the antibiotics-containing culture medium for 61 days, ten individual antibiotics-resistant clones were isolated, expanded, and analyzed by genomic PCR. Fig. 6 shows that all of the ten clones subjected to CRISPR/Cas9-induced HDR were homozygous knockouts; however, all of the ten clones subjected to the homologous recombination retained the wild type *LpMT* allele, demonstrating that they were heterozygous knockouts. Furthermore, the 803 bp PCR amplicon was absent in three of the clones (#1, 4, and 5) subjected to the homologous recombination, suggesting that the donor DNA was integrated into *LpMT* with an unexpected orientation or into another locus in the *L. passim* genome. Thus, CRISPR/Cas9–induced HDR is able to replace two alleles of an endogenous gene in *L. passim* with a single donor DNA containing an antibiotics-resistant gene.

**Figure 6.**
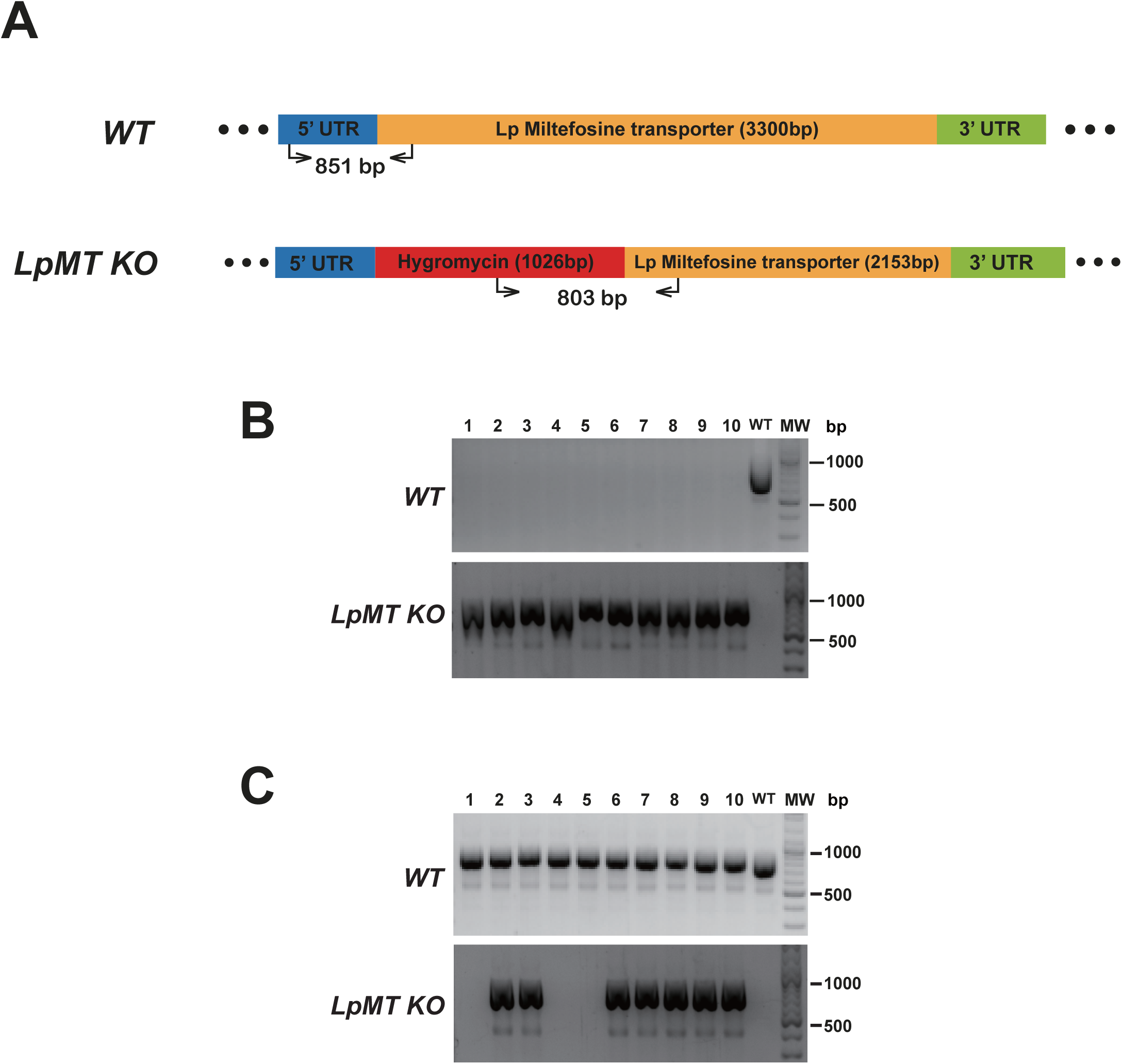
Comparison of knocking out LpMT by CRISPR/Cas9-induced HDR and homologous recombination. **(A)** Wild type (*WT*) and knocked out (*LpMT KO*) alleles of *LpMT* are shown as in Fig. 3A. The positions of primers to detect *LpMT KO* allele are shown with the expected size of PCR amplicon. **(B)** Ten antibiotics (blasticidin, G418, and hygromycin) resistant clones (1-10) isolated by CRISPR/Cas9-induced HDR together with wild type *L. passim* were analyzed by genomic PCR to detect *WT* (851 bp amplicon) and *KO* (803 bp amplicon) alleles of *LpMT*. The position of 500 and 1000 bp DNA molecular weight marker (MW) is shown at the right. **(C)** Ten hygromycin resistant clones (1-10) isolated by homologous recombination were analyzed as in (B)

## Discussion

### Stable expression of exogenous protein in *L. passim*

We successfully introduced several plasmids to express exogenous proteins (tdTomato, GFP, and Cas9) in *L. passim*. Although we evaluated the electroporation under various conditions, the efficiency of transient transfection was quite low (< 1.21 %), so that stable transfectants had to be selected by antibiotics. Although the introduced plasmid DNA does not appear to integrate into the *L. passim* genomic DNA, it is likely to be present in multicopy, since the parasite has to undergo many cell divisions to lose the plasmid DNA. Consistent with the phylogenetic similarity of *L. passim* with other trypanosomatids (Schwarz et al., 2015), the *T. cruzi* and *L. donovani* rRNA promoters are functional for protein and sgRNA expression in *L. passim*. Therefore, various expression plasmids constructed for genome editing of *Trypanosoma* and *Leishmania* by CRISPR/Cas9 could be directly applied to *L. passim* as well. *L. passim* expressing a fluorescent protein, such as GFP or tdTomato, could be useful to monitor how the parasite establishes the infection in the honey bee hindgut.

### Gene knockout in *L. passim* by CRISPR/Cas9-induced HDR

We were able to successfully knock out two endogenous genes of *L. passim* by CRISPR/Cas9-induced HDR. In contrast to knocking out a specific gene by homologous recombination, single transfection with sgRNA-expressing plasmid DNA and donor DNA containing an antibiotics-resistant gene was sufficient. However, it usually takes more than 60 days to obtain the homozygous knockout parasite by selecting the antibiotic resistant clones in the culture medium followed by isolating single clones either from agar plates or by serial dilution. Considering that we obtained both heterozygous and homozygous knockouts for *LpMT* and *LpTAT* (Figs. 3 and 4), as well as the low transfection efficiency of *L. passim*, the replacement with the donor DNA probably occurs initially with one allele. CRISPR/Cas9-induced HDR then follows with the second allele, where the first replaced allele serves as the template. This mechanism is similar to the “mutagenic chain reaction” used for converting heterozygous to homozygous mutations in fruit flies (Gantz and Bier, 2015). To shorten the selection period, electroporation of heterozygous knockout parasites with donor DNA containing different antibiotics-resistance genes followed by selection using two antibiotics should be considered. However, if the target gene is essential for the survival of *L. passim*, only heterozygous knockout clones will be selected. This was indeed the case for the paraflagellar rod component par4 (*LpPFR4*) gene, although the *PFR1* and *PFR2* genes have been successfully knocked out in *T. cruzi* (Lander et al., 2015b).

We did not observe any alterations in the target gene when we expressed only Cas9 and sgRNA in *L. passim.* After selecting *L. passim* expressing Cas9 and sgRNA using blasticidin and neomycin in the culture medium, the genomic DNA extracted from the pooled parasites or the expanded individual clones was analyzed by sequencing the PCR products encompassing the sgRNA-target site. We did not find any changes within the DNA sequences, and the same results were also obtained by digesting the PCR products with T7 endonuclease I. As previously reported (Passos-Silva et al., 2010), the NHEJ pathway is absent in trypanosomatids; however, the MMEJ pathway is apparently present in *L. donovani* and *T. cruzi* based on the successful gene modifications by expressing Cas9 and sgRNA (Peng et al., 2015; Zhang and Matlashewski, 2015). In *L. passim*, both the NHEJ and MMEJ pathways might not exist, perhaps because the parasite lacks the essential genes. Alternatively, Cas9 introduces only a single-strand break, but not DSBs in *L. passim* genomic DNA, so that only the HDR pathway is induced as a result.

We could apply the CRISPR/Cas9-induced HDR to prepare a library of *L. passim* clones in which specific genes are knocked out. The genes essential for survival could be identified by the absence of homozygous knockouts, and the genes necessary for optimal growth in the culture medium could also be tested. More importantly, we could infect honey bees with the abovementioned *L. passim* clones, and identify the genes important for establishing and maintaining the infection in the honey bee gut. Understanding the gene functions will provide insights into the molecular and cellular mechanisms of host (honey bee)-parasite (*L. passim*) interactions.

## Supporting information

## Author contribution

QL conducted all experiments. TK supervised the research project.

## Acknowledgements

We thank Jing He, Xinyi Li, and Yuntao Wang for their contribution to conduct some of the experiments. We are grateful to Dr. Duo Peng at Harvard T.H. Chan School of Public Health for suggestion on the experiments and testing *L. passim* gene annotation file with the sgRNA design website. We are also grateful to Dr. Wenwei Zhang at McGill University for suggestion on the experiments.

